# Revealing the spatial pattern of brain hemodynamic sensitivity to healthy aging through sparse DCM

**DOI:** 10.1101/2023.10.16.562585

**Authors:** Giorgia Baron, Erica Silvestri, Danilo Benozzo, Alessandro Chiuso, Alessandra Bertoldo

**Affiliations:** Department of Information Engineering, University of Padova, 35131 Padova, Italy; Padova Neuroscience Center, University of Padova, 35131 Padova, Italy

## Abstract

Age-related changes in the BOLD response could reflect neuro-vascular coupling modifications rather than simply impairments in neural functioning. In this study, we propose the use of a generative dynamic causal model (DCM) to decouple neuronal and vascular factors in the BOLD signal, with the aim of characterizing the whole-brain spatial pattern of hemodynamic sensitivity to healthy aging, as well as to test the role of hemodynamic features as independent predictors in an age-classification model.In this view, DCM was applied to the resting-state fMRI data of a cohort of 126 healthy individuals in a wide age range, providing reliable estimates of the hemodynamic response function (HRF) for each subject and each region of interest. Then, some features characterizing each HRF curve were extracted and used to fit a multivariate logistic regression model to predict the age class of each individual. Ultimately, we tested the final predictive model on an independent dataset of 338 healthy subjects selected from the Human Connectome Project Aging (HCP-A) and Development (HCP-D) cohorts. Our results entail the spatial heterogeneity of the age effects on the hemodynamic component, since its impact resulted to be strongly region- and population-specific, discouraging any space-invariant corrective procedures that attempt to correct for vascular factors when carrying out functional studies involving groups with different ages. Moreover, we demonstrated that a strong interaction exists between some specific hemodynamic features and age, further supporting the essential role of the hemodynamic factor as independent predictor of biological aging, rather than a simple confounding variable.

**Significance statement:** By inferring region-wise hemodynamic profiles at the individual level, this is the first study providing an exhaustive whole-brain characterization of the hemodynamic sensitivity to healthy aging, reporting further evidence of the vascular changes across the adult lifespan. Using a predictive framework, we analysed the statistical influence of advancing age on individual regional hemodynamic attributes, offering a quantitative evaluation of the diverse hemodynamic bias across different brain regions. We unveiled a specific set of hemodynamic predictors to discriminate young from elderly people, mainly describing vascular properties of right-hemispheric areas. This suggests the asymmetric nature of vascular degeneration processes affecting the human brain at the latest stage of life, other than a potential biomarker that could be relevant for brain-age prediction.

## Introduction

BOLD functional magnetic resonance imaging (fMRI) is widely used to study neural functioning, often overlooking its undirect nature that arises from the coupling between neuronal activity and the hemodynamic response function (HRF). Indeed, many studies have characterized BOLD-derived measures such as functional connectivity (FC), by assuming that the HRF is fixed across subjects and brain regions. This hypothesis of hemodynamic equivalence can be accepted when dealing with young healthy adults, but may not hold in older individuals (Geerligs et al., 2017), in which vascular differences can severely alter the neuro-vascular coupling. Given its critical role in defining cerebrovascular integrity (Tomoto et al., 2020), the impact of age on the vascular component of BOLD is one of the greatest sources of signal variability. As a result, a failure to consider age-related changes in the neuro-vascular coupling can induce a bias in the study of the aging brain functioning, other than a misunderstanding of its cognitive relevance (Tsvetanov, Henson, & Rowe, 2021). A fully quantification of the impact of age on the HRF is then needed since a spatial- and age-invariant canonical HRF would not accurately represent the variability induced by aging (G. Chen et al., 2023).

Among the proposed approaches to estimate the HRF, those involving the computational modelling of the fMRI signal have been widely investigated (Tsvetanov, Henson, & Rowe, 2021). While the unpredictable resting-state brain activity cannot be inferred with the traditional general linear model (Raichle & Raichle, 2001), decomposition approaches may be implemented to detect non-neuronal sources of BOLD activity (Tong et al., 2015; Tsvetanov et al., 2018). However, the selection of the spurious signals may vary depending on the chosen statistical criterion (Tsvetanov, Henson, & Rowe, 2021). Scaling methods such as the resting-state fluctuation amplitude (RSFA) have been proposed as an index of vascular reactivity, but the effects of age on RSFA cannot be fully explained by cerebral blood flow and vascular reactivity, making it dangerous to employ it as a normalization signal (Tsvetanov, Henson, Jones, et al., 2021). Additionally, many works have employed different deconvolution methods (Caballero-Gaudes et al., 2019; G. Chen et al., 2023; Karahanoǧlu et al., 2013; Wu et al., 2013). While some techniques evaluate the BOLD signal against a given threshold to identify resting pseudo-events, possibly introducing detection errors which tend to bias the recovery of HRF shape (Wu et al., 2013), others employ a fixed HRF shape throughout the brain (Karahanoǧlu et al., 2013), limiting the description of the high variability of HRF within and across subjects.

Differently from the previous approaches, DCM implements a Bayesian inference procedure to effectively disentangle the neuronal and vascular parameters from the fMRI signal associated with each brain region (Daunizeau et al., 2011; Friston et al., 2003).

Very few studies have investigated the relevance of DCM-based HRF in neurocognitive aging focusing on specific regions and/or vascular features (Anderson et al., 2020; Tsvetanov et al., 2016). Here we propose the use of a sparse version of DCM (Prando et al., 2020) to fully characterize the HRF at the subject level and separately for each region of interest. We then extracted a set of vascular features from each HRF curve (West et al., 2019) to explore their spatial variability in two groups of young and old participants, thus providing a whole-brain quantitative evaluation of the diverse hemodynamic sensitivity to healthy aging. We finally examined the hypothesis that certain hemodynamic features hold significance in predicting age by training an age-classification model and testing it on an independent second dataset of healthy subjects.

As far as we know, this is the first study to provide a quantitative evaluation of the spatial pattern of the hemodynamic sensitivity to advancing age, as well as to give a whole-brain overview of the role of the hemodynamic component as an independent predictor of biological aging, rather than a simple normalization or confounding variable.

## Materials

We analysed resting-state functional MRI data in 464 normal adults between the ages of 18-100 years from two independent cohorts. The first dataset was retrieved from the publicly available MPI-Leipzig Mind-Brain-Body dataset (MPI-LMBB) (Babayan et al., 2019), while the second data collection includes subjects recruited from the Human Connectome Project in Aging (HCP-Aging) and Development (HCP-Development) (Harms et al., 2018). In the present study, the MPI-LMBB was employed as the primary dataset for model derivation and internal validation. The combination of data collected from HCP-Aging and HCP-Development served as the testing dataset used to assess the generalizability of the derived predictive model.

### Participants

#### Dataset 1

The first resting-state fMRI dataset employed in this study consists of resting-state scans of 295 healthy subjects from the publicly available MPI-Leipzig Mind-Brain-Body dataset (https://legacy.openfmri.org/dataset/ds000221/). The data selection was performed on the original dataset (consisting of 318 individuals) by excluding participants with high motion (less than 400 volumes with a mean framewise displacement (FD) <0.4 mm) or affected by pre-processing failures and/or unavailability of rs-fMRI data (Moretto et al., 2022). Then, while the first half of the dataset was employed for clustering purposes (see detailed description in *Data acquisition and preprocessing* section), a fraction of the second half was used for the setup of the model after discarding middle-aged participants (31-59 years of age). After discarding subjects for which the algorithm could not converge to a solution in a reasonable amount of time, a final sample of 95 younger (20-30 yo) participants and 31 older (60-80 yo) participants were included in the study.

#### Dataset 2

HCP-Aging and HCP-Development are multisite projects designed to generate an in-depth imaging, behavioural and biosample repository of typical brain changes across a broad portion of the human lifespan (Harms et al., 2018), with HCP-D recruiting participants from 5 to 21 years of age and HCP-A enrolling individuals from 36 to >100 years of age. Both datasets used in this study was drawn from the second release (Lifespan HCP 2.0 Data Release) and focused on typical brains only since HCP studies excluded participants who have diagnosed and treated for major neuropsychiatric, neurological disorders or depression, as well as those affected by cognitive impairment, learning disabilities or severe medical conditions (Harms et al., 2018). We did not consider young subjects collected from the former HCP-Young Adult study (Van Essen et al., 2012), since this dataset differs in some important scanning protocol choices from HCP-A/HCP-D that can entail differences in the downstream measures derived from MRI (Harms et al., 2018). From both datasets, we extracted data from 641 subjects aged 18-21 and 36-100 years from HCP-D and HCP-A respectively, in order to approximately match the age range associated with the training dataset (NDA study DOI: http://dx.doi.org/10.15154/hk8k-pm84). As for the MPI-LMBB dataset, subjects affected by high head motion (less than 400 volumes with mean FD<0.4 mm) were discarded, as well as those subjects for which DCM did not converge to a final solution. As a result, data from 338 individuals (180 young 18-40 y and 158 old 60-100 y participants) were available for the testing stage.

For both datasets, demographic details are described in table 1.

**Table 1.** Participant demographics and Repetition Time (TR) in the considered datasets.

|  | *Dataset1* | *Dataset2* |
| --- | --- | --- |
| *Source dataset* | MPI-LMBB | HCP-A/HCP-D |
| *Young N (after sDCM)* | 95 | 180 |
| *Old N (after sDCM)* | 31 | 158 |
| *Young Age (mean/SD)* | 25.1/2.5 | 25.4/8.2 |
| *Old Age (mean/SD)* | 68.2/4.6 | 73.6/9.6 |
| *Young gender (M/F)* | 78/17 | 86/94 |
| *Old gender (M/F)* | 17/14 | 79/79 |
| *TR (s)* | 1.4 | 0.8 |

### Data acquisition and preprocessing

#### Dataset 1

Data acquisition was performed with a 3T Siemens Magnetom Verio scanner, equipped with a 32-channel head coil. The protocol included a T1-weighted 3D magnetization-prepared 2 rapid acquisition gradient echoes (3D-MP2RAGE) and resting-state fMRI images acquired using a 2D multiband gradient-recalled echo echo-planar imaging (EPI) sequence (TR = 1400 ms; flip angle = 69°; voxel size = 2.3 × 2.3 × 2.3 mm; volumes = 657; multiband factor = 4) and two spin echo acquisitions. Further details about the acquisition protocol can be found in (Babayan et al., 2019). During rs-fMRI scans, the subjects were asked to keep their eyes opened and to lie down as still as possible.

The structural preprocessing pipeline included bias field correction (*N4BiasFieldCorrection* (Tustison et al., 2010)), skull-stripping (*MASS* (Doshi et al., 2013)) and nonlinear diffeomorphic registration (Avants et al., 2011) to the symmetric MNI152 2009c atlas (Fonov et al., 2011). Pre-processing of rs-fMRI data consisted of slice timing (Smith et al., 2004), distortion *topup* (Andersson et al., 2003) and motion correction *mcflirt* (Jenkinson et al., 2002)) and nonlinear normalization to the symmetric MNI atlas (Fonov et al., 2011) passing through the single subject pseudo-T1w image, which was obtained by multiplying the T1w 3D-MP2RAGE image with its second inversion time image (boundary-based registration (Greve & Fischl, 2009)). As a second step, the GIFT toolbox (http://trendscenter.org/software/gift/) was used to decompose the functional pre-processed data into independent components (ICs) by performing a group spatial-ICA. The ICs were manually classified by visual inspection of both the spatial maps and the source power spectra, in accordance with (Damaraju et al., 2014; Griffanti et al., 2014). As a result, ICs that were related to banding artifacts, vascular or noise components were discarded. Further details on the IC selection can be gathered in (Silvestri et al., 2022). Then, 10 principal components related to CSF and white matter signal (5 from WM, 5 from CSF) were regressed out from rsfMRI timeseries as well as the 6 standard head motion parameters and their temporal derivatives (Jo et al., 2013).

#### Dataset 2

Neuroimaging data of HCP-A/D was collected with a Siemens 3 Tesla Prisma system with a 32-channel head coil. Anatomical T1-weighted images were acquired via a multiecho magnetization-prepared gradient-echo sequence, while resting-state fMRI images were acquired using a 2D multiband gradient-recalled echo EPI sequence (TR = 800 ms; flip angle = 52°; voxel size = 2.0 × 2.0 × 2.0 mm; volumes = 478; multiband factor = 8). Two sessions of eyes-open resting state fMRI (REST1 and REST2) were performed with opposite phase-encoding direction (four different runs). Additionally, a pair of opposite phase-encoding spin-echo were acquired and used to correct functional images for signal distortion. Further details about the acquisition protocol can be found in (Harms et al., 2018).

Concerning data preprocessing, the data package employed in the study is the preprocessed volumetric version provided by HCP (Glasser et al., 2013), which contains cleaned rfMRI files aligned across subjects using nonlinear volume registration. The dataset includes outputs of HCP Functional Preprocessing for resting state scans, which is the result of applying Generic fMRI Volume Processing Pipeline, linear detrending and *multirun* FIX (MR-ICA-FIX) implemented in the HCP pipelines (Glasser et al., 2018). For each subject, only the first run (encoded as *rfMRI_REST1_AP_hp0_clean*) was employed in the following analysis to ensure phase-encoding consistency with *Dataset 1*.

### Common preprocessing steps

#### Application of the functional atlas

For both datasets, we extracted time series data from a 100-area parcellation scheme of the cortex provided by the Schaefer atlas (Schaefer et al., 2018), which maps to 7 resting-state functional networks (RSNs) referring to both hemispheres: Visual network (Vis) (5 parcels), Somatomotor network (SomMot) (6 parcels), Dorsal attention network (DorsAttn) (9 parcels), Saliency/Ventral attention network (SalVenAttn) (11 parcels), Limbic network (Limbic) (5 parcels), Control network (Cont) (10 parcels), Default mode network (DMN/Default) (16 parcels).

We also defined a set of 10 subcortical regions based on the AAL2 segmentation (Rolls et al., 2015). For each hemisphere, we selected 5 regions consisting of thalamus proper, caudate, putamen, pallidum and hippocampus.

We then assigned to each subject a binary temporal mask accounting for brain volumes corrupted by head motion (FD>0.4 mm) and we applied volume despiking to the time series by means of the *icatb_despike_tc* function of the GIFT toolbox. Moreover, the temporal traces were band-pass filtered (0.0078 to 0.2 Hz) so as to retain the range of frequencies that is usually supported by the canonical HRF in the DCM framework (J. E. Chen & Glover, 2015; Henson & Friston, 2007).

#### Consensus clustering

Given the need to keep the computational load of sparse DCM at a reasonable level (Prando et al., 2020), a consensus clustering procedure was applied to reduce the number of cortical parcels derived from the Schaefer atlas, and at the same time to keep the hemodynamic analysis in a whole brain setting. As a result, we applied to the first half of the MPI-LMBB dataset (147 subjects) a Consensus Clustering Evidence Accumulation (CC-EA) procedure, similarly to what described in (Ryali et al., 2015), to determine the number of optimal clusters in rs-fMRI time series data. Specifically, this framework employs base and consensus clustering methods to get robust and stable clusters. First, various groupings are obtained for each individual dataset using base clustering method; then, consensus clustering is applied on these groupings to get stable clusters across individuals. In order to account for hemodynamic differences across spatially distant parcels, this procedure was performed selectively for subsets of adjacent cortical regions referring to the same functional network. This additional constraint ensured that only functionally homogeneous and spatially contiguous parcels could be grouped together. For sets composed of only 2 contiguous regions of interest (ROIs), the objective criteria for determining the optimal number of clusters could not be applied. In these cases, the grouping has been performed if the pair Pearson’s correlation was on average greater than the mean correlation within the other clusters. This approach ensures that the hemodynamic consistency of each cluster is preserved, contributing to reflect the whole-brain vascular complexity (G. Chen et al., 2023; Taylor et al., 2018) underlying variations of regulatory neurochemical processes (Rangaprakash et al., 2018). The resulting clustering procedure provided 62 cortical clusters as displayed in Table 1, from which demeaned fMRI time courses (i.e., within-cluster mean BOLD signal) were extracted and supplied as inputs to sparse DCM together with the BOLD signals from subcortical sources.

## Methods

Figure 1 provides a schematic representation of the whole pipeline.

**Figure 1.**
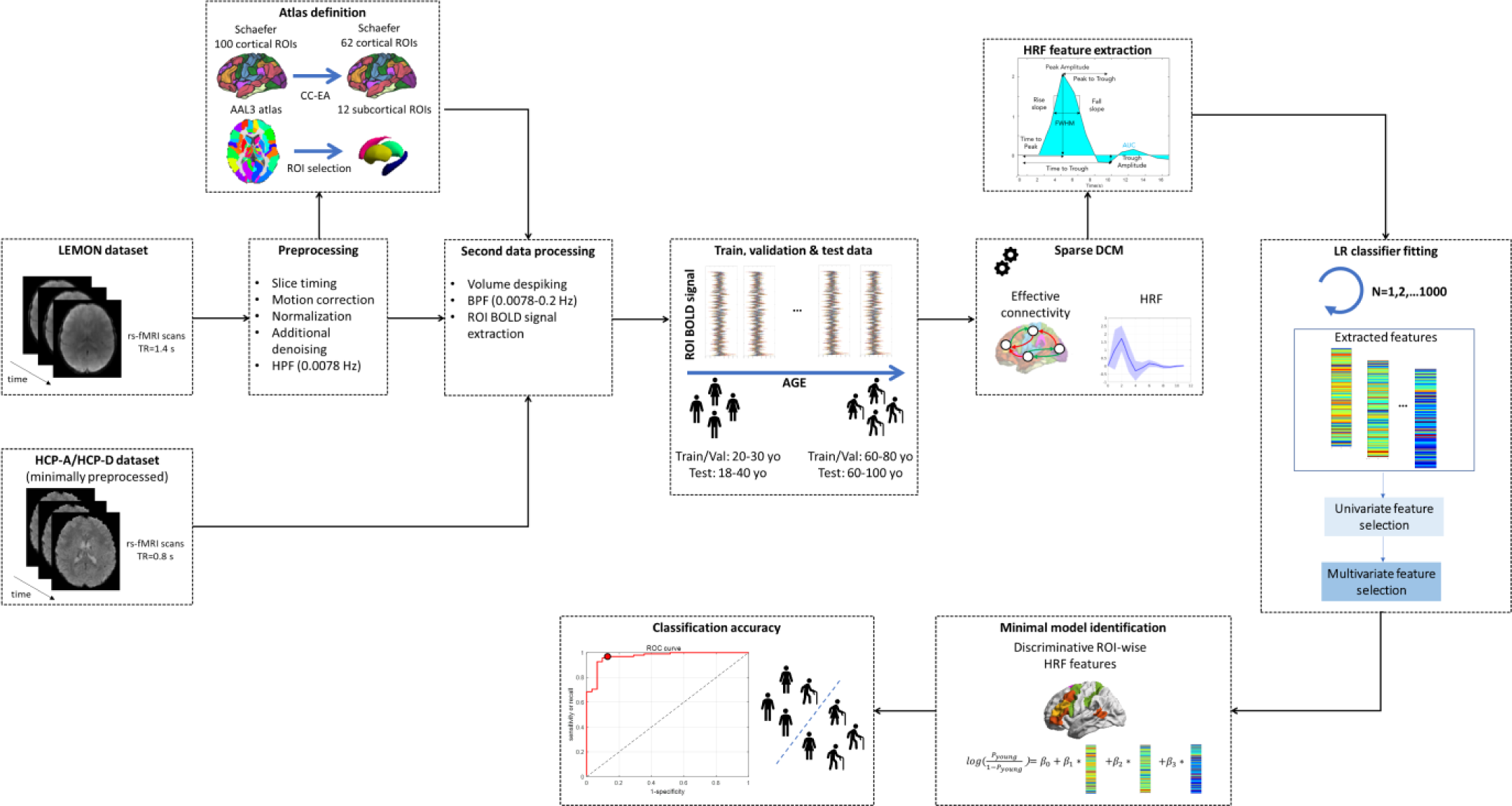
Overview of the key processing steps for the implementation of the age-classification model to assess the predictive power of sDCM-derived hemodynamic features in healthy aging. HPF: high-pass filter, BPF: band-pass filter, yo: years old.

### Hemodynamic inference

#### Sparse DCM

For each subject, the resting-state hemodynamic responses and the effective connectivity matrix were inferred using the sparse DCM framework (Prando et al., 2020). More specifically, thanks to a discretization and statistical linearization procedure, the inversion of such a linear generative model allows to estimate subject-level parameters accounting for the hemodynamic variability across brain areas, differently from other DCM approaches holding a fixed HRF(Handwerker et al., 2004). In this view, intra-subject haemodynamic variability is captured by the formal modelling of the neuro-vascular coupling process through differential equations that mathematically describe the link of the hemodynamic and neuronal connectivity parameters with the measured fMRI signal. More in detail, the hemodynamic linearization and discretization procedure consists in translating empirical priors on the physiological parameters describing the nonlinear model of the hemodynamic response (Buxton et al., 1998) into a hemodynamic prior which is exploited in the estimation process of the HRF. This is motivated by the low temporal resolution of fMRI data, which usually ranges from 0.5 to 3 seconds, and by the idea that the hemodynamic response can be modelled as a Finite Impulse Response (FIR) model with the neuronal state as input and the BOLD signal as output. In order to ensure that the length of the input response was large enough to model relevant temporal dependencies, we set the hemodynamic length equal to 16.8 s for both datasets, verifying that the number of samples was sufficient to capture the region-wise HRF dynamics. Concerning connectivity parameters, this DCM variant implements a sparsity inducing mechanism that automatically prunes irrelevant connections, thereby avoiding the need to perform a selection between competing network structures (Prando et al., 2020). The algorithm has been further adjusted to account for the signal reliability of the temporal frames, by introducing the binary temporal mask as a weighting measure during the estimation procedure. It follows that the application of sDCM resulted in the estimation of the effective connectivity (EC) matrix and 72 ROI-specific HRF profiles (referring to 62 cortical and 10 subcortical nodes) at the individual level, encoding the spatial variations of the neurovascular coupling.

#### Extraction of the hemodynamic features

First, we applied a principal component analysis (PCA) on the all the estimated HRF curves to provide a coarse overview of the hemodynamic variability captured by sDCM across all subjects and brain regions. Then, as carried out in (West et al., 2019), nine features defining the HRF shape and timing properties were computed from each individual ROI-level HRF (see Figure 1), including peak amplitude (*A peak*), time-to-peak (*t peak*), rise slope, trough amplitude (*A trough*), time-to-trough (*t trough*), fall slope, peak-to-trough, full width half maximum of the peak (*FWHM*) and area under the curve (*AUC*). It follows that 648 variables (9 features for 72 ROIs) were considered in the subsequent fitting procedure.

#### Age-classification model

In order to assess the prediction power of the hemodynamic features, we trained a logistic regression model classifying young-age vs. elderly subjects. The reason for choosing this classification task between middle-age and elderly individuals relies on the aim to assess whether DCM-derived brain hemodynamic characterization (in terms of its associated features) is able to capture the effect of aging, other than quantitatively evaluating the spatial pattern of the hemodynamic sensitivity to it. The choice of training the model on the dataset with lower temporal resolution comes from the purpose of investigating hemodynamic factors that could be detectable from most commonly available MRI scanners. Then, testing the model on HCP data collected with a higher sampling rate allowed to assess its potential applicability to a wider range of fMRI acquisition and pre-processing pipelines.

#### Partitioning of the dataset

To overcome the risk of overfitting the predictive model on a predefined partition of the training dataset and to reduce the sensitivity of the model to outliers, the fitting procedure has been iterated over 1000 random partitions. Specifically, in each repetition we adopted a 20% hold-out validation approach to split the dataset 1 into two parts, training on one while validating on the other with the trained model. Given the unbalanced number of the two age classes, we performed a stratified split to maintain the class proportion in the two sets. The Pearson’s correlation coefficient of each training covariate with age has also been assessed to later consider it as a quality metrics of the given data partition in the fitting of the final logistic regression model. Then, the outputs of the single models have been combined and statistically evaluated to define a final logistic regression model. All subsequent data analysis was conducted employing code written in MATLAB R2021b (Mathworks, USA).

#### Univariate feature selection

Given the high number of identified variables, the first step regarded the application of a univariate significance test for selecting features that significantly vary across the two age groups derived from the training subset. For each of the 648 hemodynamic features, we conducted a two-sample t-test by considering the two samples as extracted from two normal populations with unequal variances (e.g., for the parcel LHVis1, we statistically compared the distributions of each hemodynamic feature referring to the two populations). A Holm-Bonferroni correction was applied to control for multiple comparisons (corrected p-value<.05). As a result, only features that show significant age differences were considered in the following fitting procedure.

#### Multivariate feature selection

We then performed a correlation analysis to reduce the collinearity effect between independent variables. Specifically, we evaluated the degree of Pearson’s correlation between each pair of hemodynamic features that were retained in the previous selection stage, and we removed very high redundancies (ρ > 0.85) by retaining only the variable that correlated the most with the output.

Subsequently, a multivariate selection of features has been performed on the training set. To increase the statistical robustness of the final selection, a bootstrap resampling was executed 100 times to create multiple internal training sets. At each iteration, we generated the internal dataset by sampling with replacement from the initial training dataset a collection of data with the same cardinality and by preserving the original weights of the two classes. Then a stepwise backward feature selection was applied to each sampled normalized internal set by using the Bayesian Information Criterion (BIC) as the stopping rule.

#### Fit of the logit model

The hemodynamic covariates that were selected in a significant portion (>50%) of the internal training sets in the previous stage have then been considered to fit a logistic linear model predicting the probability of a subject to be classified as young (p_young_). The model predictive performance has been evaluated on the validation set by computing the corresponding concordance index (*c-index*), which is numerically equivalent to the area under the Receiver Operating Characteristic curve (*AUC_ROC_*).

#### Minimal model identification

Having estimated a distinct classifier for each random partition, we combined single results to determine the hemodynamic features that were most frequently selected in the 1000 iterations. First, we filtered out atypical model realizations by assessing the data representativeness of each training subset, which was quantified by computing the Euclidean distance between the correlation vectors of the training and the whole dataset with the output variable. We further excluded any model that performed worse than a random classifier on the test subset (*AUC_ROC_* or *c-index*<0.5). We finally reduced the number of plausible variables by discarding selected features whose corresponding coefficient estimate was not statistically significant, according to a *p-value* > 0.05. This resulted in considering only features selected by the best-fitted models, which have been ordered in descending fashion based on their selection frequency. On the basis of the identification of the knee point in the frequency curve, the most frequently selected variables have then been employed to define the final logistic regression classifier. The final model predicting the age group was then retrained on the 1000 random sets to get the median regression coefficient for each hemodynamic feature, after selecting only good-fitted model realizations (*AUC_ROC_* > 0.5, *p-value* < 0.05).

#### Statistical evaluation of classification performance

The final step was then to assess the performance of the minimal model on the testing dataset (dataset 2). This evaluation was performed by computing the final ROC curve and the associated AUC (*AUC_ROC_*) to provide an aggregate measure of performance across multiple classification thresholds. The Precision Recall (PR) curve and the associated AUC (*AUC_PRC_*) was also considered since it can be more informative than the ROC plot when evaluating classifiers on imbalanced datasets. The performance of the model was finally tested by employing the value maximizing the overall balanced accuracy as the cut-off for the final prediction.

Similarly, the same metrics were assessed on each internal validation set and on the whole *Dataset 1* to provide sample confidence intervals for the performance in the random train-test split and an upper bound for the overall accuracy, respectively.

## Data and Code Accessibility

This study is a secondary analysis of publicly available datasets. The download links for the datasets are provided in the manuscript. No personal identifiable information was used in this study. Authors can make the code used for the analysis available to any reader directly upon reasonable request.

## Results

### Region-wise hemodynamic response function

Using PCA decomposition, we first reported evidence of the variability of the hemodynamic response functions inferred through sparse DCM across all subjects and brain regions. Figure 2 illustrates the distribution of HRFs as described by the first 3 principal components (PCs) in each of the two age samples (*Dataset 1*). Although the spatial localisation of clusters in terms of functional networks is quite reproducible in both groups, it is worth noticing that ROI scores differ significantly across the brain space and subjects on the basis on their hemodynamic features, providing clear evidence of the high variability characterizing HRF. As illustrated in the central panels, the hemodynamic profiles not only vary across aging groups, but they are also strictly dependent on the localisation of the brain area under investigation (i.e., inversion of the amplitude difference when moving from a cortical to a subcortical parcel), motivating the following ROI-wise analysis of the hemodynamic curves.

**Figure 2.**
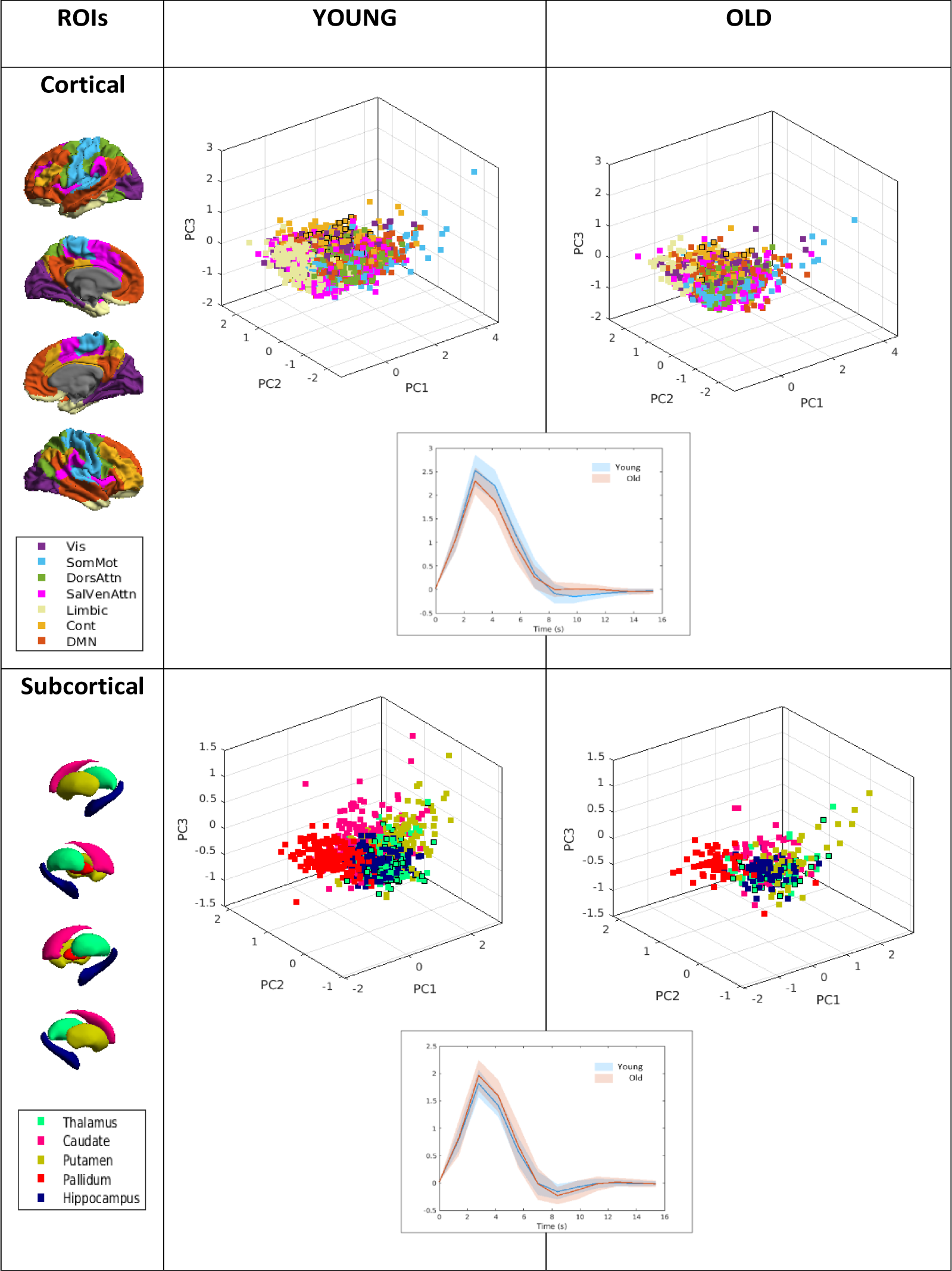
The three-dimension space identified by the first 3 PCs allows to describe the HRF variability across subjects in the two age samples and across ROIs. Each point condenses the hemodynamic description of an individual parcel. The colour denotes network affiliation or the corresponding subcortical node. Points marked with a black outline refer to an example ROI whose mean (solid line) and standard deviation (shading) of the HRF curves are plotted in the central panels, corresponding to young (blue) and old (red) healthy adults in *Dataset 1*. Three-dimensional hemispheric and subcortical renderings in this figure and in the following ones were obtained by using the ENIGMA toolbox (Larivière et al., 2021).

### Spatial and group variability of hemodynamic features

Figure 3 displays the resulting median maps of hemodynamic features characterizing the two age groups. Both cortical and subcortical maps exhibit spatial heterogeneity. Specifically, the median peak amplitude varies substantially across ROIs, with higher values in visual and frontoparietal areas, as well in caudate and pallidum subcortical nodes. However, this feature shows a general decreasing trend with aging which involves all brain regions. Similarly, trough amplitude (i.e., the post-stimulus undershoot) is quite variable across ROIs with a spatial pattern that is inversely proportional to that associated with peak amplitude in both cortical and subcortical regions, meaning that a higher hemodynamic peak is often related to a lower (i.e., deeper) post-stimulus undershoot. Concerning *AUC*, we found higher values in cortical visual and some SomMot areas as well, whose associated pattern highly resembles the one characterizing the peak amplitude. In addition, *AUC* shows high values in the caudate nucleus and putamen, while it remains low in hippocampus and thalamus. These spatial profiles are mostly preserved in the older subjects, even if characterized by a slight decrease in strength. Together with peak amplitude, this feature entails a pronounced hemispheric asymmetry (slightly declining with age) described by a higher spatial heterogeneity regarding the right side of the brain. Regarding rate features (fall and rise slope), we found similar cortical maps (whose pattern recalls the one revealed by *AUC*) but opposite in sign, with an increasing (decreasing) trend for fall (rise) slope in the group of elders. Interestingly, this trend is reversed for subcortical nodes. Concerning response width (*FWHM*), the young sample shows a broader *FWHM* in frontoparietal areas and in the putamen and caudate nuclei, while the lowest values are encountered in the hippocampus and limbic areas in both groups.

**Figure 3.**
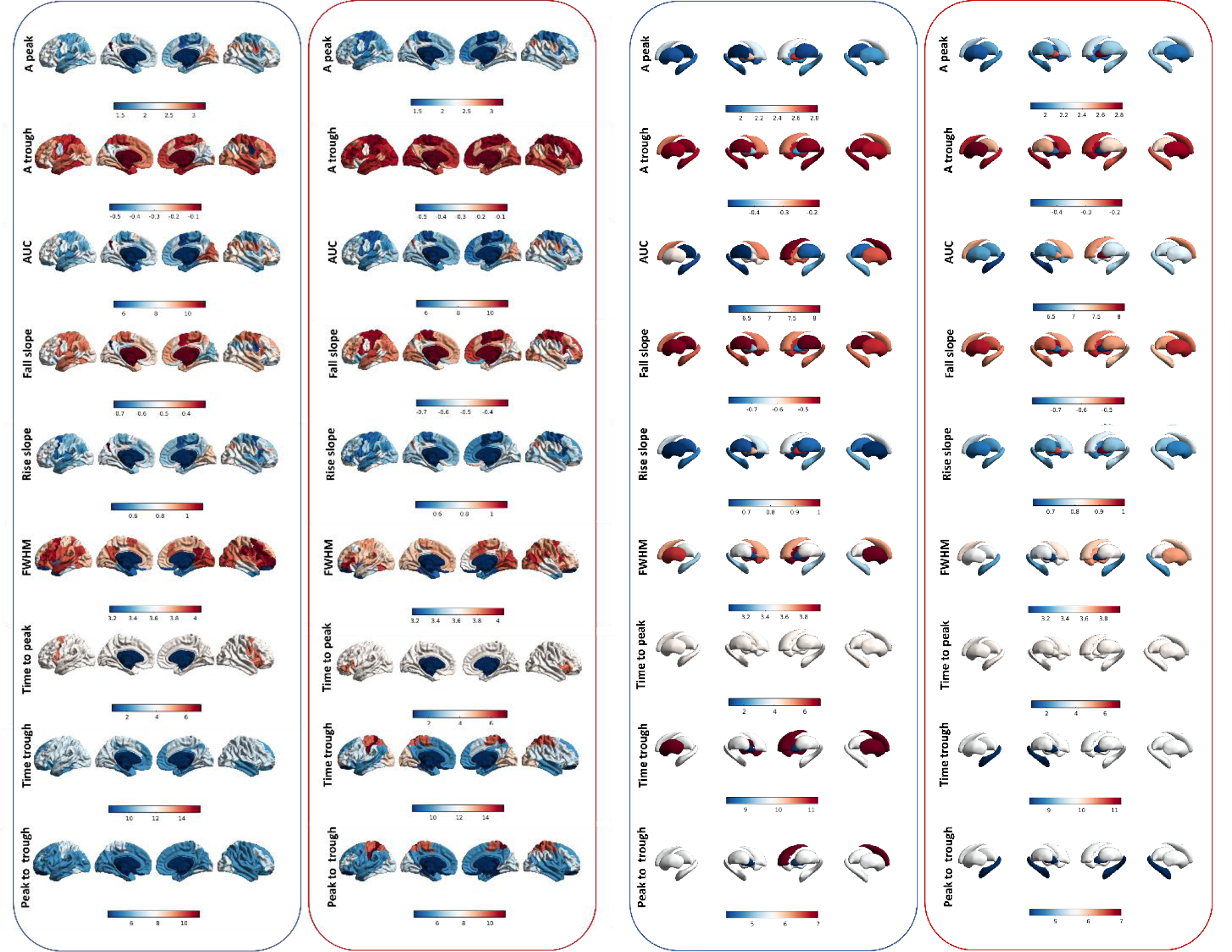
Median spatial distribution maps for the HRF features computed as described in Figure 1. Blue boxes refer to the young sample, while red boxes are related to the old sample (*Dataset 1*). Different scales are used for cortical and subcortical regions, while the same scale is adopted for each feature and surface to ease the comparison between the age groups.

On the other hand, timing features (i.e., time-to-peak, time-to-trough, peak-to-trough) generally show a weaker variability than the previous hemodynamic features. Time-to-peak varies very little across the cortex and is stable across subcortical nodes, revealing the reduced sensitivity of this feature to age-related changes. In the older subjects, time-to-trough shows a pronounced increase in specific SomMot and DorsAttn areas in both hemispheres. Concerning subcortical areas, we observed the highest values in the right caudate nucleus and putamen in the younger group, while the lowest values characterize the left hippocampus and pallidum in the elders. Regarding peak-to-trough, its cortical median map exhibits a very similar cortical pattern. However, a slightly different spatial scheme is encountered in subcortical nuclei, where the right caudate presents the highest value in the younger sample, while hippocampus and pallidum in older participants are interested by the lowest peak-to-trough.

### Univariate feature selection

The univariate analysis involving HRF differences between younger and older groups across the 1000 iterations yielded distinct results depending on the selected ROI and the feature of interest (Figure 4). Specifically, significant differences were mostly observed in areas of lower-order functional networks (i.e., visual and somatosensory-motor), as well as in more cognitive RSNs such as DorsAttn, Cont and DMN. Selection frequency is rather hemispheric balanced, although the most selected HRF feature may vary based on the considered ROI. However, it is worth noticing that amplitude (i.e., peak amplitude, trough amplitude and *AUC*) and rate features (fall and rise slope) generally prevail over the others in terms of sensitivity to age-related hemodynamic changes. Concerning subcortical nodes, thalamic areas are the most frequently selected for significant differences in peak amplitude and fall slope. For each variable with a selection frequency > 0, Figure 4b reports the corresponding values of difference between the group means. While peak amplitude, *AUC* and rise slope are generally greater in younger, undershoot amplitude is usually damped (i.e., higher values) in elders, as well as the fall slope. Note that for these features the general trend is reversed in subcortical areas. On the contrary, *FWHM* differences are consistent in sign across the brain space. As expected, temporal features show very weak differences (mainly values prevailing in the old group) across nodes, which are mainly localized in Vis, DorsAttn and some subcortical parcels.

**Figure 4.**
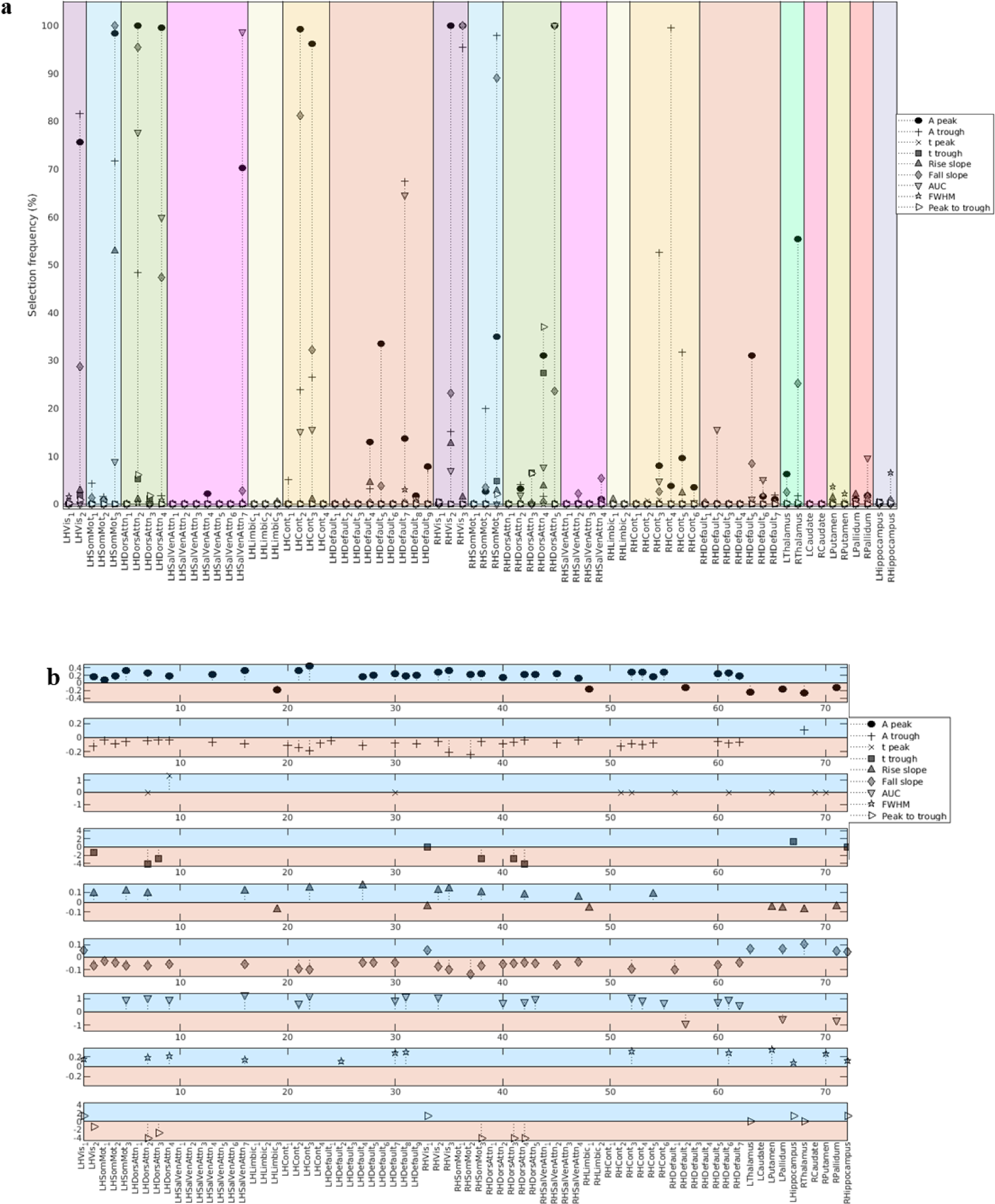
Results of the univariate feature selection step. (a) Frequency plot illustrating the percentage of frequency selection for each feature across the 1000 dataset partitions, reporting a balanced trend across hemispheres. Each marker refers to a different hemodynamic feature (as indicated in the legend), while the colour code discriminates between functional networks and/or subcortical ROIs. (b) For each feature-ROI pair characterized by a selection frequency > 0 in panel a, the stem plot reports the difference between the corresponding median values in the two age samples as extracted from the maps in Figure 3. Positive values indicate a significantly higher estimate in the young group (red area), while negative values refer to a significantly higher estimate in the old group (blue area).

### Multivariate feature selection

As illustrated in Figure 5a, a totally different scenario is provided after the multivariate selection process. First, the results suggest a prominent hemispheric asymmetry, since retained hemodynamic features are mostly localized in the right-hand side of the brain for both cortical and subcortical domains. This points out that right-hemisphere ROIs generally perform better in discriminating the age class than the left counterpart in terms of their corresponding vascular properties. Specifically, the most predicting variables are found in Vis, SomMot, DorsAttn, Cont and DMN networks. Concerning subcortical nuclei, right thalamus and pallidum are the only nodes that play an important role in the model prediction. As regards feature class, amplitude features prevail after multivariate selection, corroborating their crucial contribution in predicting age. Figure 5b outlines the top 10 HRF features in descending order according to their multivariate selection frequency, together with the corresponding univariate selection frequency computed in the previous step. It is evident that the two selection stages follow two completely different trends, motivating the importance of accounting for multivariate interactions of hemodynamic features for model fitting. Specifically, we observed an evident slope change in the multivariate selection curve (i.e., knee point), which we employed to identify the 5 HRF features to be included in the final minimal logit model.

**Figure 5.**
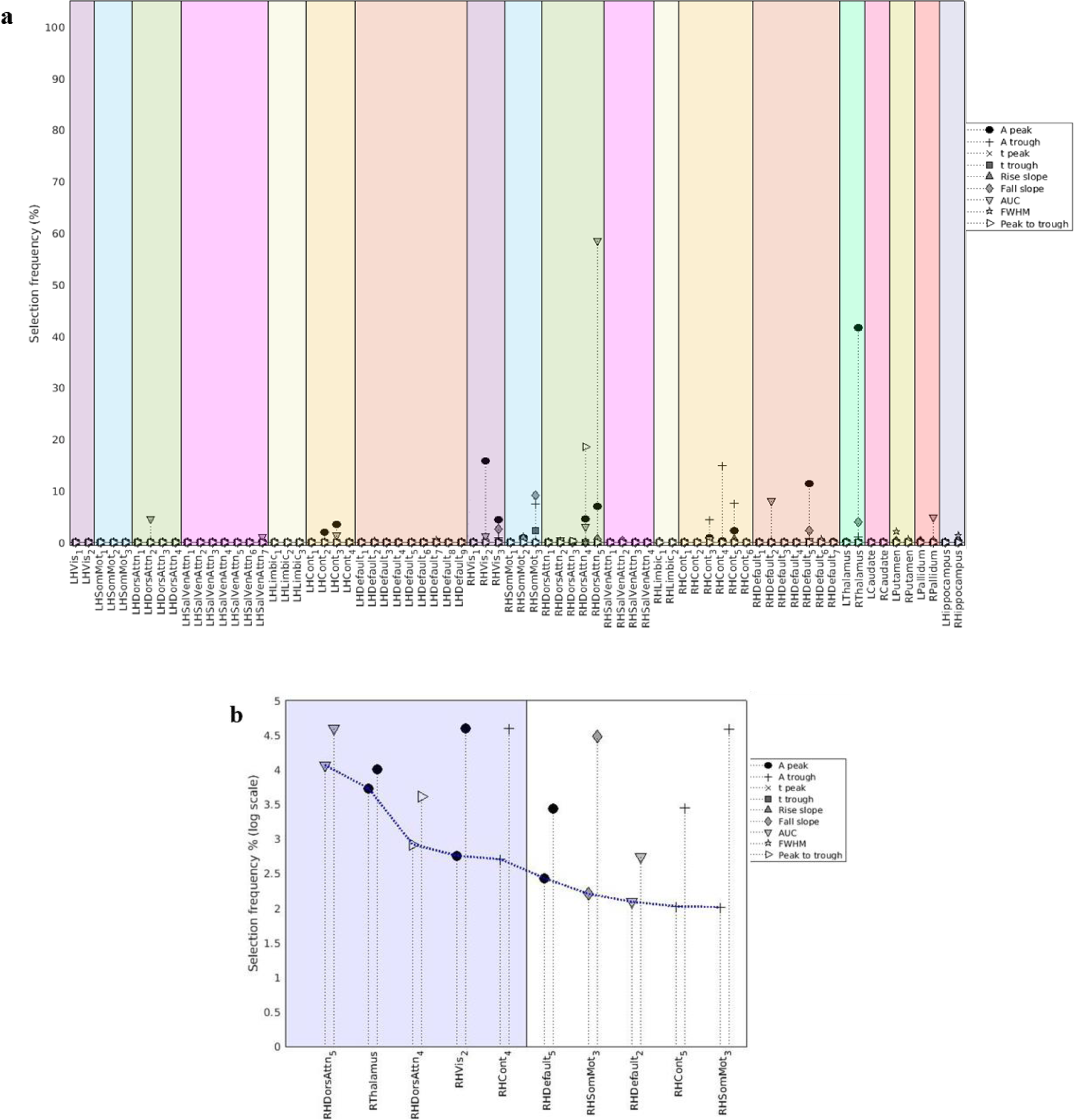
Results of the multivariate feature selection step. (a) Frequency plot illustrating the percentage of frequency selection for each feature across the 1000 dataset partitions, reporting a lateralized trend in favour of right-hemispheric areas. Each marker refers to a different hemodynamic feature (as indicated in the legend), while the colour code discriminates between functional networks and/or subcortical ROIs. (b) Stem plot reporting the multivariate (first stem) and univariate (second stem) selection frequency values for each of the top 10 hemodynamic features (in descending order). The blue area highlights the features that have been included in the final minimal predictive model (see Figure 6).

### Minimal model identification

Figure 6a depicts the distributions of regression coefficients obtained by fitting the minimal model on the 1000 random sets. Despite the presence of outliers, the width of the boxplots is quite limited, suggesting a good consistency of estimated β values in terms of variance across model realizations. Regarding cortical parcels, the 4 selected ROIs are mostly localized in the right frontoparietal cortex (involving dorsal attention *PrCv, FEF* (*DorsAttn5*), *Post5* (*DorsAttn4*) and control *PFCl3* areas) and in a great part of the right visual central/temporal-occipital region. On the other side, right thalamic nucleus is the only subcortical node implicated in the model prediction. To what concerns HRF feature class, we can deduce that the age groups can be discriminated on the basis of the *AUC* in the first dorsal attention ROI (positive relation with logit of *p_young_*), peak amplitude of the right thalamus and the visual parcel (negative and positive relation with logit of *p_young_*, respectively), peak to trough for the second dorsal attention ROI (negative relation with logit of *p_young_*), trough amplitude in the case of the control network region (negative relation with logit of *p_young_*). On the other hand, Figure 6b displays the estimated PDFs for performance metrics of the 1000 model realizations.

**Figure 6.**
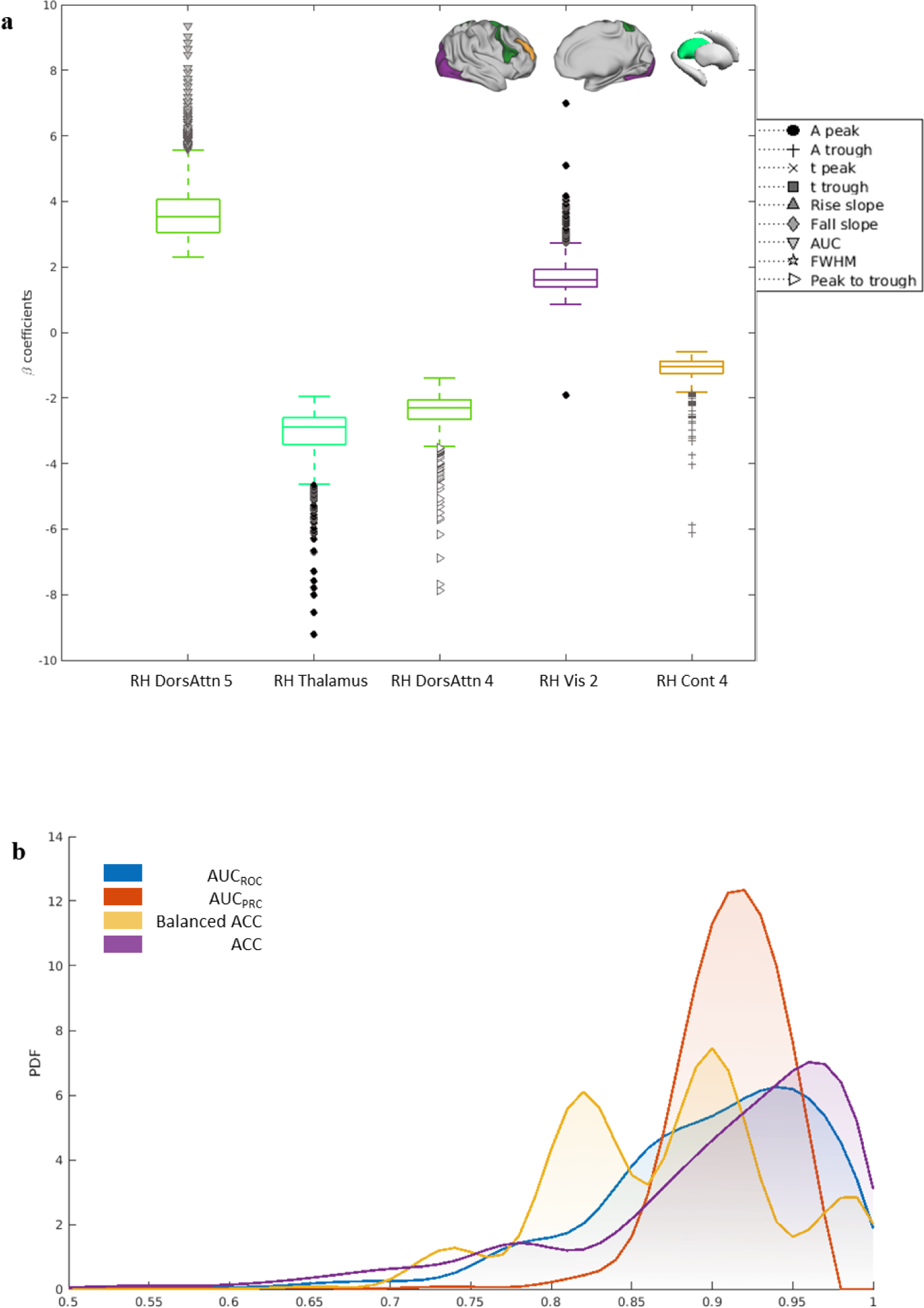
Results of the minimal model identification. (a) Distributions of the regression coefficients obtained by fitting the minimal model across the 1000 set realizations. Each marker refers to a different hemodynamic feature (as indicated in the legend), while the colour of the boxplots discriminates between functional networks and/or subcortical ROIs. Spatial localization of the corresponding ROIs (right hemisphere) is provided by the brain surface renderings on the top. (b) Probability density functions (PDFs) of the performance metrics (*AUC_ROC_*, *AUC_PRC_*, balanced accuracy and overall accuracy) obtained by evaluating the minimal model realizations across all 1000 random sets.

Overall, the results suggest that the minimal model performs well across all the set partitions in terms of (balanced) accuracy, as well as for precision, specificity, and sensitivity (i.e., *AUC_ROC_* and *AUC_PRC_*), since all distributions mostly range from 0.7 to 1.

### Classification performance of the minimal model

Having derived the median values of regression coefficients depicted in Figure 6a, we tested the “median” minimal model both on the whole *Dataset 1* and *Dataset 2*. Panels a and b in Figure 7 display the ROCs and PRCs in both datasets, respectively. The receiver operating characteristic and precision recall curves indicate good classification performance for both datasets (i.e., better than the random classifier), although PRC trend generally exhibits a higher variability in the HCP datasets. Coloured dots overlaid on the curves denote the cut-off scores maximizing the balanced accuracy (specifically, 0.92 in *Dataset 1* and 0.79 in *Dataset 2*), whose corresponding value has been employed to identify the confusion matrices in Figure 7c and 7d. The asymmetry of the confusion chart in panel c suggest that the classification error is due to some young subjects being classified as old, while all elders are correctly classified by the minimal model. The predictive scenario is more balanced in *Dataset 2*, despite the percentage of right predictions is slightly higher in the old cohort.

**Figure 7.**
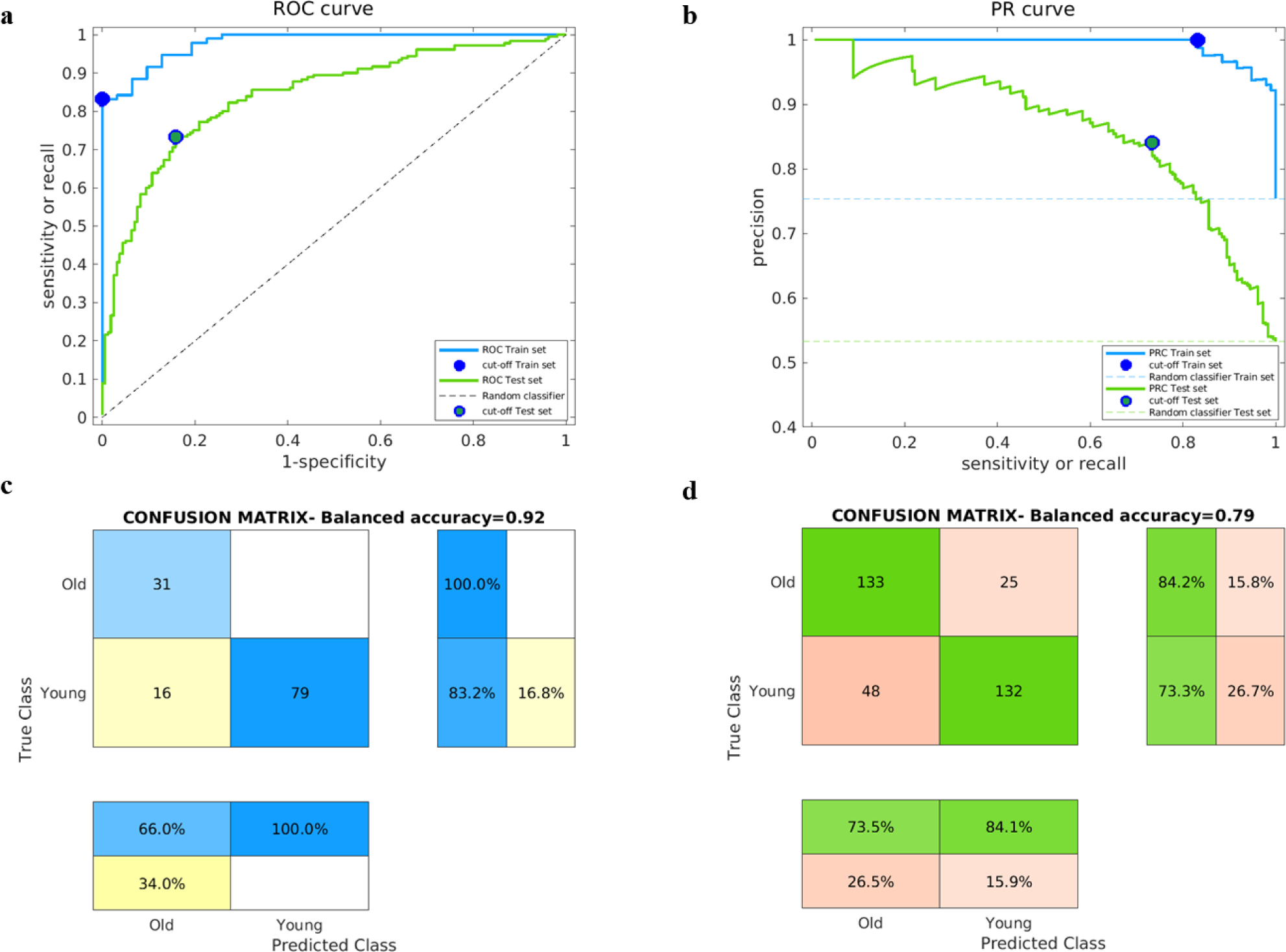
Classification performance of the minimal model. (a) Confusion matrix of the minimal model for the prediction of the age class in *Dataset 1*; for each aging group, the class-wise precisions and class-wise recalls (computed as percentage of the number of correctly and incorrectly classified observations for each predicted and true class, respectively) are indicated. (b) Confusion matrix of the minimal model for the prediction of the age class in *Dataset 2*; for each aging group, the class-wise precisions and class-wise recalls (computed as percentage of the number of correctly and incorrectly classified observations for each predicted and true class, respectively) are indicated. (c) ROCs of the minimal model for the prediction in *Dataset 1* (blue) and *Dataset 2* (green), with coloured dots denoting the cut-off values for *Dataset 1* (blue) and *Dataset 2* (green); the dashed line corresponds to the ROC curve of the random classifier. (d) PRCs of the minimal model for the prediction in *Dataset 1* (blue) and *Dataset 2* (green), with coloured dots denoting the cut-off values for *Dataset 1* (blue) and *Dataset 2* (green); the dashed line corresponds to the PR curve (i.e. the positive rate P/(P+N)) of the random classifier.

## Discussion

### Heterogeneity of aging effects on HRF

First, the PCA analysis reported evidence of the high inter-subject and regional variability of the hemodynamic response (Handwerker et al., 2004; Kassinopoulos & Mitsis, 2019; Rangaprakash et al., 2018), supporting the need, for each subject, to model HRF locally in the brain. Given the reported modifications of the hemodynamic system with healthy aging (Ances et al., 2009; Y. Li et al., 2018; West et al., 2019), we analysed how the hemodynamic profile varies in space over different brain areas and between the two age groups by providing an exhaustive description of the median spatial pattern of each feature and testing for significant hemodynamic differences between the two populations through the univariate selection step. Assessing the spatial pattern of these features can provide additional insights into the age-related effects of the underlying neurovascular factors such as CBF or CMRO_2_, which are known to specifically affect HRF amplitude, undershoot and time-to-peak (Arichi et al., 2012; G. Chen et al., 2023; Kim & Ress, 2016).

In general, amplitude features shared correlated trends across the lifespan, with significant age-related decreases for peak height (Ances et al., 2009; Fabiani et al., 2014; Hutchison et al., 2013; Ward et al., 2015; West et al., 2019) and *AUC*, which are reflected in a damping of post-stimulus undershoot in the same networks. This is line with (West et al., 2019), which reports the same spatial trends for peak and trough amplitude, while it finds an inverse (not significant) tendency for *AUC*. As expected, spatial variability of rate features varies with aging by a decreasing (increasing) trend for rise (fall) slope, accordingly to the results illustrated in (West et al., 2019). The emergence of such aging HRF properties possibly relates to a significantly slower neurovascular coupling in elderly people (Cherkaoui et al., 2021). In this view, also timing features generally present an increasing age-related trend in cortical ROIs, which is consistent to the results presented in (West et al., 2019).

Interestingly, the parametric patterns of significant age-related changes are generally reversed in subcortical areas, where the HRF is usually reported to be significantly faster and narrower than in cortical ROIs (Kim et al., 2022; Lewis et al., 2018). This is in line with (Lee et al., 2009), which represented prior findings of augmented subcortical perfusion in healthy older adults. Specifically, aging seems to have a greater impact on thalamic HRF, which presents significant differences in terms of a greater peak amplitude and faster rise and fall phases. This contrasts with (Galiano et al., 2020), which reported that perfusion reduces with age in the medial thalamus. Also, right pallidum, whose *AUC* appears to significantly increase with age, is the only subcortical nucleus to exhibit a significant decreasing impact on the association between age and CBF in (Y. Chen et al., 2011). This lack of concordance between results could derive from the low signal-to-noise ratio of the fMRI signal in these areas (Goodro et al., 2012; Singh et al., 2018) and the impact of the volume reduction observed in normal aging (Wang et al., 2019).

### The prominent role of right-hemispheric regions in the age-classification task

Since univariate age-related differences are largely confined to the local level without integration at the whole-brain level, we then assessed the combined prediction power of the proposed HRF features that were retained in the univariate selection step, in order to classify young vs. elderly subjects using a standard machine learning tool. The reason for performing such classification task proceeds from the aim to assess whether the hemodynamic component could reflect brain aging independently from neural information, motivating its potential role as biomarker of brain age. Moreover, this supervised multivariate learning step allows to rank the most discriminative features by also considering interacting effects between hemodynamic variables. Previous studies have already reported evidence of the predictive power of HRF in healthy aging by examining a limited number of cortical nodes and networks (Anderson et al., 2020; Tsvetanov et al., 2016) and/or a reduced set of hemodynamic features describing the deviation from a canonical HRF shape (Cherkaoui et al., 2021). On the contrary, the proposed predictive framework provides a whole-brain spatial characterization of the hemodynamic sensitivity to healthy aging, by fully describing age-related vascular alterations of the HRF shape in terms of amplitude, rate and timing features. Our results show that right-hemispheric ROIs generally have an improved prediction power to discriminate age compared to their left counterpart, unveiling the prominent role of inter-hemispheric asymmetry in neurovascular coupling. This result is in line with aging theories speculating the validity of a right-hemi aging model (Dolcos et al., 2002) and other studies reporting hemispheric asymmetries in functional activity with increasing age (Beason-Held et al., 2008; Z. Li et al., 2009; Zuo et al., 2010). Basing on our observations, these results could be biased by the pronounced hemodynamic alterations affecting the rightward ROIs, thus reflecting asymmetric neurodegenerative processes of vascular nature rather than physiological neural plasticity (Cherkaoui et al., 2021).Moreover, the minimal model allowed to identify which brain regions are more predictable of the age class. Concerning cortical networks, some Cont and DorsAttn areas appear to be vulnerable to hemodynamic decline in aging. This might lead to reconsider previous works reporting decreased within-DorsAttn connectivity and cohesion with increasing age (Betzel et al., 2014; Zonneveld et al., 2019), further suggesting the potential bias in these results induced by the hemodynamic component. Likewise, hemodynamic differences emerge in the Cont network, which was shown to have a general within-network decrease of local activity in older adults (Bethlehem et al., 2020; Geerligs et al., 2015). Interestingly, (Avelar-Pereira et al., 2017) examined the interactions between the right Cont and DorsAttn, reporting that older adults show altered connectivity values compared to the young, while connectivity between the left Cont and DorsAttn was not significantly different between the age groups. Specifically, the significant age-sensitive HRF detected in the right *PFCl* recalls the results provided by (Grady et al., 2016), which reported that this region shows stronger connections to the DorsAttn in older adults. Similarly, the peak amplitude detected in right visual central areas and right thalamus seems to be a crucial predictor of brain age. This observation is consistent with the study of (Beason-Held et al., 2008), which points out a decreased CBF in occipital and occipitotemporal cortices that may impact visual perception in older subjects. Moreover, the same study illustrated a longitudinal increase of CBF in the right-hemispheric subcortical thalamus, proposing that this trend may reflect preservation of function over time in view of age-related declines in global cerebral blood flow.

Overall, the minimal model exhibits outstanding predictive performance in both *Dataset 1* and *Dataset 2*, reaching an overall balanced accuracy of 92% and 79%, respectively, without losing in precision. Although model fitting was performed on the lower temporal resolution dataset, the minimal model predicts HCP subjects’ age class with a good accuracy. As a result, the pattern of unveiled hemodynamic predictors could potentially represent a reliable biomarker for brain-age that could be replicable for a wide range of fMRI acquisition protocols.

## Limitations

As previously stated, fMRI temporal resolution significantly matters for accurate timing and rate feature estimation (Rangaprakash et al., 2018), whose reliability in our study is limited by the high acquisition repetition time. Especially for subcortical nuclei, HRF assessment is critical because the quantification requires high spatiotemporal resolution to resolve the small nuclear subdivisions of subcortical brain regions (Singh et al., 2018). Moreover, the proposed regional analysis is conditioned by the choice of the parcellation atlas. It would be interesting to deepen this research by testing the reproducibility of the predictive hemodynamic pattern with some atlas variations (Cherkaoui et al., 2021). Third, we did not include effective connectivity parameters as candidate predictors of age (Tsvetanov et al., 2016), since our aim was to test the independent predictive ability of vascular factors. However, including connectivity factors into the predictive model could provide a more complete interpretation of the unique and shared contributions of neuronal and vascular components to brain age. Additionally, future studies might be strengthened by handling the higher number of predictors through deep learning methods, so that unknown relationships between variables can be more accurately identified (Cole et al., 2017).

## Conclusions

By inferring ROI-wise hemodynamic profile at the individual level through sDCM, we provided a whole-brain characterization of the spatial pattern of brain hemodynamic sensitivity to healthy aging. Through the univariate testing, we evaluated the statistical impact of increasing age on each regional hemodynamic feature, proposing a quantitative assessment of the spatially heterogeneous hemodynamic bias that could affect the functional behaviour of cortical regions and the main subcortical nuclei. We then unveiled a specific set of hemodynamic predictors to discriminate young from elderly people, mainly describing vascular properties of right-hemispheric areas devoted to cognitive functions. This suggests the asymmetric nature of vascular degeneration processes affecting the aging human brain, other than a potential candidate biomarker underlying the ‘age’ of the brain to make individualised predictions about neurovascular diseases and mortality risk in older adults (Green & Hillersdal, 2021).

## Acknowledgments

Giorgia Baron and Danilo Benozzo were supported by the DEI Proactive grant “Personalized whole brain models for neuroscience: inference and validation” from the Department of Information Engineering of the University of Padova (Italy).

The HCP-Aging dataset reported in this study was supported by grants of the National Institute On Aging of the National Institutes of Health under Award Number U01AG052564 and by funds provided by the McDonnell Center for Systems Neuroscience at Washington University in St. Louis. The HCP-Aging 2.0 Release data used in this report came from DOI: 10.15154/1520707<http://dx.doi.org/10.15154/1520707>.

The HCP-Development dataset reported in this study was supported by grants of the National Institute Of Mental Health of the National Institutes of Health under Award Number U01MH109589 and by funds provided by the McDonnell Center for Systems Neuroscience at Washington University in St. Louis. The HCP-Development 2.0 Release data used in this report came from DOI: 10.15154/1520708<http://dx.doi.org/10.15154/1520708>.

